# MBPpred: Proteome-wide detection of membrane lipid-binding proteins using profile Hidden Markov Models

**DOI:** 10.1101/034942

**Authors:** Katerina C. Nastou, Georgios N. Tsaousis, Nikolaos C. Papandreou, Stavros J. Hamodrakas

## Abstract

A large number of modular domains that exhibit specific lipid binding properties are present in many membrane proteins involved in trafficking and signal transduction. These domains are present in either eukaryotic peripheral membrane or transmembrane proteins and are responsible for the non-covalent interactions of these proteins with membrane lipids. Here we report a profile Hidden Markov Model based method capable of detecting Membrane Binding Proteins (MBPs) from information encoded in their amino acid sequence, called MBPpred. The method identifies MBPs that contain one or more of the Membrane Binding Domains (MBDs) that have been described to date, and further classifies these proteins based on their position in respect to the membrane, either as peripheral or transmembrane. MBPpred is available online at http://bioinformatics.biol.uoa.gr/MBPpred. This method was applied in selected eukaryotic proteomes, in order to examine the characteristics they exhibit in various eukaryotic kingdoms and phylums.

## 1. INTRODUCTION

A cell’s structure and functions rely significantly on membranes, since they are responsible for its compartmentalization and are associated with nearly half of all its proteins [1]. Membrane proteins are of central importance as they take part in a large variety of cellular functions such as ion, metabolite and macromolecular transport and signal transduction [2], as well as cell adhesion, cell-cell communication, protein anchoring to specific locations in the cell, control of membrane lipid composition and the organization and maintenance of organelle and cell shape [3, 4]. These proteins can either be embedded directly within the lipid bilayer (transmembrane proteins), or can be associated with the membrane indirectly via interactions with membrane proteins or lipids (peripheral membrane and lipid-anchored proteins) [5]. Transmembrane proteins constitute ~20 to 30% of fully sequenced proteomes [6] and they are the most studied class of membrane proteins. Consequently, many prediction methods have been designed specifically for this class of proteins through the years and have been improved and optimized using several different implementations [7].

Peripheral membrane proteins interact non-covalently with the membrane, either directly via membrane lipids or indirectly with transmembrane proteins. Directly interacting membrane proteins usually have domains that allow for the specific or non-specific interaction with membrane lipids [8]. Besides peripheral membrane proteins, these domains are also present in extramembranous regions of transmembrane proteins [9] – either intracellular or extracellular – and are known as Membrane Binding Domains (MBDs). MBDs are of great importance to the cell, since proteins that contain such domains take part in a variety of cellular processes such as cell signaling and membrane trafficking, vital for the cell’s survival and growth. While MBDs of the PH superfamily have recently been found in prokaryotic proteins [10], the main focus of experimental studies is on eukaryotic membrane binding domains and representatives of other membrane binding proteins are restricted mainly in eukaryotes [11]. Homologs of such “eukaryote-specific” MBDs can be discovered in prokaryotes with genome-wide approaches, even though their function might differ from that of their eukaryotic counterparts. Computational studies that indicate the existence of domains in prokaryotes that act as membrane binding have been conducted, and particularly domains like BON [12] and Nisin [13] have been characterized as putative membrane-binding domains. However, the lack of experimental evidence regarding these domains in the organisms in which they are found is a stumbling block towards discovering their function.

MBDs are extremely diverse and their only common characteristic is their non-covalent interaction with membrane lipids, with different affinities. A significant number of MBDs have been identified to date. While some of them, like C2, and BAR [14] have been extensively studied in the last decades, mainly with experimental methods, there is a growing number of recently identified MBDs for which very little is known, such as IMD and GOLPH3 [15]. Structural studies have aided in the elucidation of the interactions of MBDs with the membrane. However, the search of new membrane binding domains with experimental methods would be immensely time-consuming and expensive. Thus, the development of genome-wide prediction methods for the detection of membrane binding proteins is necessary.

A large number of Membrane Binding Proteins (MBPs) act as enzymes by recognizing specific lipid head groups. Mutations of these proteins affect their molecular function, and a number of diseases have been described, that are attributed to the malfunction of these proteins [16]. Despite their importance, and the fact that there have been extensive structural studies regarding these proteins [14], MBPs have not been studied comprehensively with computational methods. Only two methods that allow for the detection of peripheral proteins from the existence of such domains have been reported to date. The first method, developed in 2006 [17], was based on structural characteristics of these proteins and the second, developed in 2010 [18], on information encoded in amino acid sequence. However, neither one of these methods is currently available online.

The comprehension of the molecular mechanisms that Membrane Binding Proteins use to perform their functions will be extremely significant for the unraveling of their activity inside cells. The augmentation of large scale proteomic and computational studies of Membrane Binding Domains and proteins harboring them, will aid immensely towards achieving this goal in the next few years.

We report here the design and development of a sequence-based method that identifies Membrane Binding Proteins in eukaryotic proteomes with the use of profile Hidden Markov Models (pHMMs), specific to membrane binding domains (MBDs). The method also classifies Membrane Binding Proteins (MBPs) according to their relationship with the membrane, and thus allows for the detection of peripheral membrane proteins.

## 2. METHODS

After an extensive literature search 18 domains were identified (Annexin, ANTH, BAR, C1, C2, ENTH, Discoidin, FERM, FYVE, Gla, GOLPH3, GRAM, IMD, KA1, PH, PX, PTB, Tubby) for which well-established biochemical and crystallographic experimental data for the interaction with membrane lipids exist. Each of these domains was mapped to at least one characteristic pHMM from the Pfam database [19], since in our case the majority of these profiles are well defined. Subsequently, a pHMM library (MBDslib) containing 40 pHMMs that were derived from Pfam was created. The mapping between the different pHMMs and the 18 MBDs is shown in Table 1.

**Table 1.** Mapping between the pHMMs of MBDslib and known MBDs. In the first column of this table the Membrane Binding Domains that were isolated from literature are shown and in the second column the unique pHMM identifier from the Pfam database.

| MBD | pHMMs |
| --- | --- |
| Annexin | PF00191 |
| ANTH | PF07651 |
| BAR | PF03114, PF10455, PF16746 |
| C1 | PF00130, PF03107, PF07649 |
| C2 | PF00168, PF14429, PF00792, PF10409 |
| Discoidin | PF00754 |
| ENTH | PF01417 |
| FYVE | PF01363, PF02318 |
| Gla | PF00594 |
| IMD | PF08397 |
| KA1 | PF02149 |
| PH | PF00169, PF08458, PF14593, PF15404, PF15405, PF15406, PF15409, PF15410, PF15411, PF15413, PF16457, PF16652, PF14844 |
| PTB | PF08416 |
| GOLPH3 | PF05719 |
| PX | PF00787 |
| Tubby | PF01167 |
| GRAM | PF02893 |
| FERM | PF00373, PF09380, PF09379 |

The MBPpred algorithm consists of two levels: a detection and a classification level. To develop the detection level of MBPpred, the HMMER package was utilized in order to search the pHMM library MBDslib, and detect Membrane Binding Proteins (MBPs) from a set of protein sequences. Using the hmmsearch program of HMMER, one can “search” the aforementioned library, and thus identify proteins which belong to the families used to create the library and, subsequently, find MBPs from a set of protein sequences. The classification level of MBPpred was created, in order to distinguish MBPs into transmembrane and peripheral membrane proteins with the use of the PredClass algorithm [6]. PredClass classifies proteins into four distinct classes, namely membrane, globular, mixed and fibrous. Proteins, in the first class are actually only transmembrane proteins, while MBPs in the second and third class are considered peripheral MBPs.

MBPpred was evaluated with cross-validation in order to assess its performance. In specific, a jackknife cross-validation experiment was conducted, to validate the pHMMs performance for the detection of sequences not used in the creation of the profiles (pseudo-novel sequences). For each one of the 40 profiles in the library, one sequence was removed from the multiple sequence alignment (MSA) of the pHMM’s seed set, and a new pHMM was constructed from the remaining sequences in the MSA. Then the profile’s ability to correctly classify the removed sequence, as well as the 500 sequences from the negative dataset was measured [20].

In addition, two datasets that were assembled from PDB [21] were used to evaluate MBPpred and the method was also compared with the predictor, which was developed by Bhardwaj et al. [17], in 2006.

In order to create non-redundant sequence datasets BLASTClust [22] was used, both for the creation of the positive and the negative datasets. Protein sequences with less than 30% sequence identity with each other, in a sequence length coverage of 90%, were retrieved using this program. The positive dataset consists of known MBPs, which target the membrane via MBDs. Initially it consisted of 202 proteins, 71 of which were non-redundant. After the removal of proteins present in the seed sequence sets of the pHMMs used to create MBDslib, the final positive dataset was assembled, which consists of 49 non-redundant proteins. The negative dataset was retrieved from a PDB search for eukaryotic proteins that do not have membrane or lipid binding properties, as described in their PDB files and contained 9057 nonredundant sequences. 500 sequences were randomly chosen from this dataset, in an attempt to balance the negative and positive datasets, while maintaining the information needed to evaluate our method (Table S1). In order to compare MBPpred with the predictor developed by Bhardwaj et al. [17] more precisely, the two datasets introduced in that study were used, one of membrane and one of non-membrane binding proteins, both with known three-dimensional structures. The negative dataset consists of 225 proteins and the positive of 35 proteins (Table S1). It should be noted, that this method used only 9 (ANTH, C1, C2, ENTH, FYVE, PH, PX, Tubby, BAR) out of the 18 MBDs incorporated in MBPpred.

For the prediction performance of MBPpred five measures were used, namely Accuracy, Sensitivity, Specificity, Balanced Accuracy and Matthew’s Correlation Coefficient. True/false positives (TP, FP) and true/false negatives (TN, FN) were counted on a per protein basis.

Accuracy is the proximity of measurement results to the true value and is calculated as:

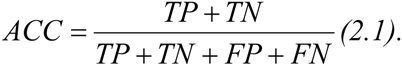

Sensitivity, or true positive rate is:

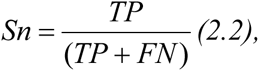

and Specificity, or true negative rate is:

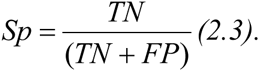

Besides these measures, the balanced accuracy and Matthew’s Correlation Coefficient (MCC) were used to appraise the performance of MBPpred. Balanced accuracy is the average of sensitivity and specificity and, together with MCC, is considered a better measure [23] when the data sizes of the positive and negative datasets are not balanced. MCC is calculated as:

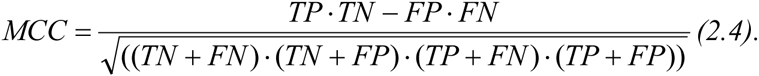

Moreover, MBPpred was applied to 30 selected (Table S2) and all 407 reference eukaryotic proteomes (Table S3) retrieved from UniProtKB [24] (release: 2015_12) in order to identify potential membrane binding proteins that interact with lipids non-covalently and to perform a quantification analysis regarding these proteins. MBPpred was also applied in all Bacterial and Archaean reference proteomes from UniProtKB.

## 3. RESULTS AND DISCUSSION

### 3.1. The MBPpred Algorithm

The detection level of MBPpred uses a library of 40 pHMMs, which correspond to 18 Membrane Binding Domains (MBDs) that were identified from literature. This library is used for the detection of Membrane Binding Proteins (MBPs). If, during a search of the library with HMMER, the score of an alignment between a query protein and at least one of the profiles is higher than the gathering threshold of each pHMM (as reported in Pfam), then the protein is characterized as a MBP. An analysis was performed, where different scoring thresholds than those defined by Pfam, were used. This analysis showed that, when tested against the proteins of the evaluation dataset, best results were retrieved with the use of the gathering thresholds and not with other more or less strict cut-offs (Table S4). Proteins that score higher than the threshold for at least one of the domains, in the library, are characterized as possible membrane binding.

The classification level of MBPpred uses the PredClass algorithm in order to classify MBPs, in respect to their interaction with the membrane, into peripheral or transmembrane. PredClass’s speed and the use of information solely encoded by amino acid sequences makes this algorithm suitable for the implementation of the classification level of our algorithm.

**Figure 1.**
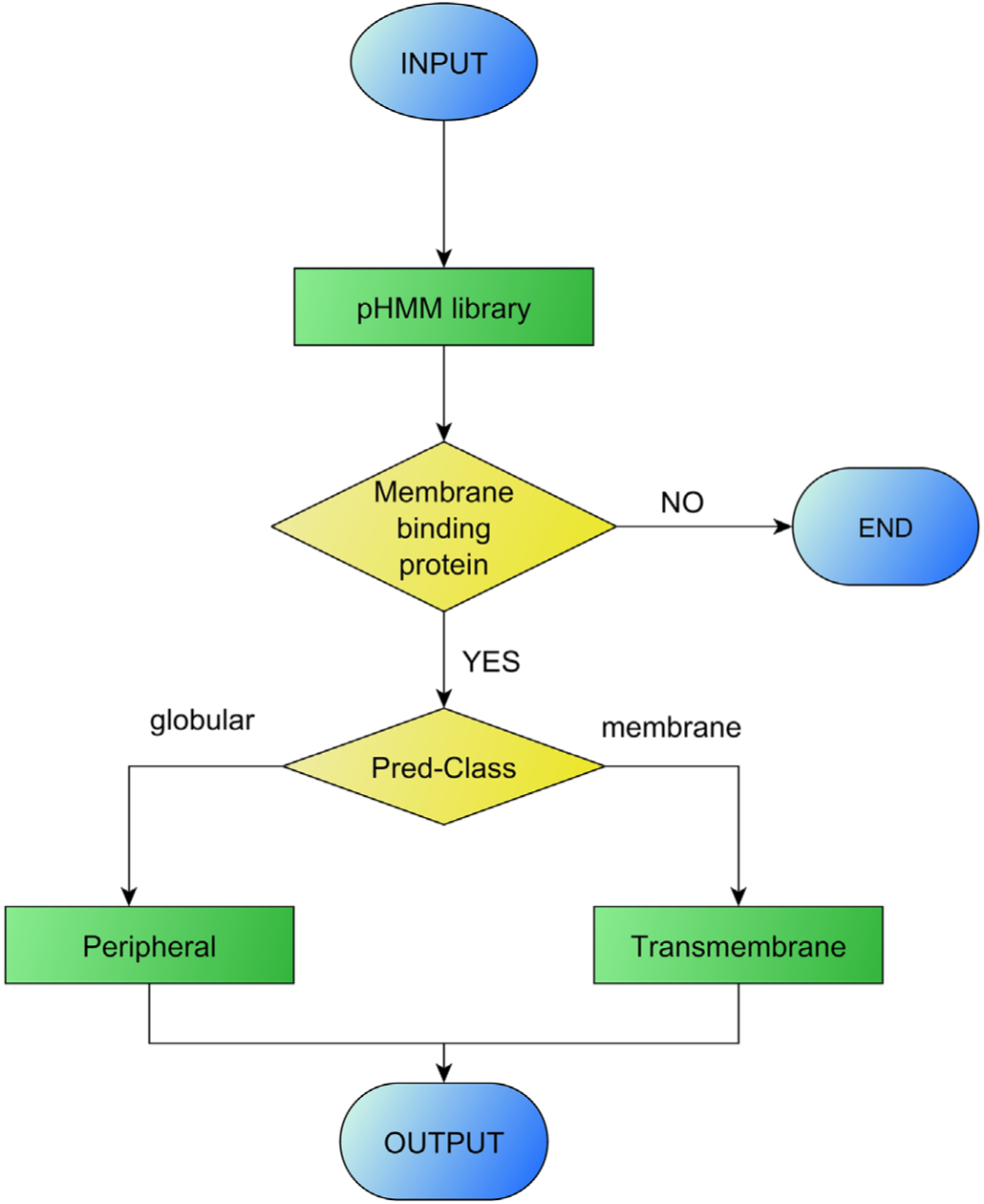
Flowchart of the MBPpred algorithm

### 3.2. Web interface of MBPpred

A web interface has been developed for MBP-Pred and the method is publicly available through http://bioinformatics.biol.uoa.gr/MBPpred. Through the main page, the user can access the query submission page, the manual and contact pages. Query submission can be performed by either pasting a single or a set of protein sequences in the textbox provided, or by uploading a file with fasta formatted sequences. Even though the method is meant to be used and has been tested with eukaryotic proteins, a user can also use the method on prokaryotic sequences in the search of homologous sequences to those of eukaryotic MBPs; results from such submissions should be carefully interpreted regarding the function of these proteins, since the role of prokaryotic proteins with domains that act as membrane lipid-binding in eukaryotes has not been unveiled yet [10, 11, 25] and different, unknown to date, mechanisms could be involved in the function of these proteins in other domains of life that could result from the radically different cell and membrane structures of these organisms.

After a successful query submission users are transferred to the results page where they can gather information about their submission, as well as extensive information in downloadable files about MBPs (if any). The final results files contain a protein identifier, the position and score of the domain(s) present in the protein and the type of membrane protein (peripheral or transmembrane) along with its sequence and length. The output files produced by MBPpred and their contents are shown in Fig. 2. MBPpred is fast, since for a query length the size of the human proteome the algorithm produces results in ca. five minutes, which makes it sufficient for proteomic scale applications.

**Figure 2.**
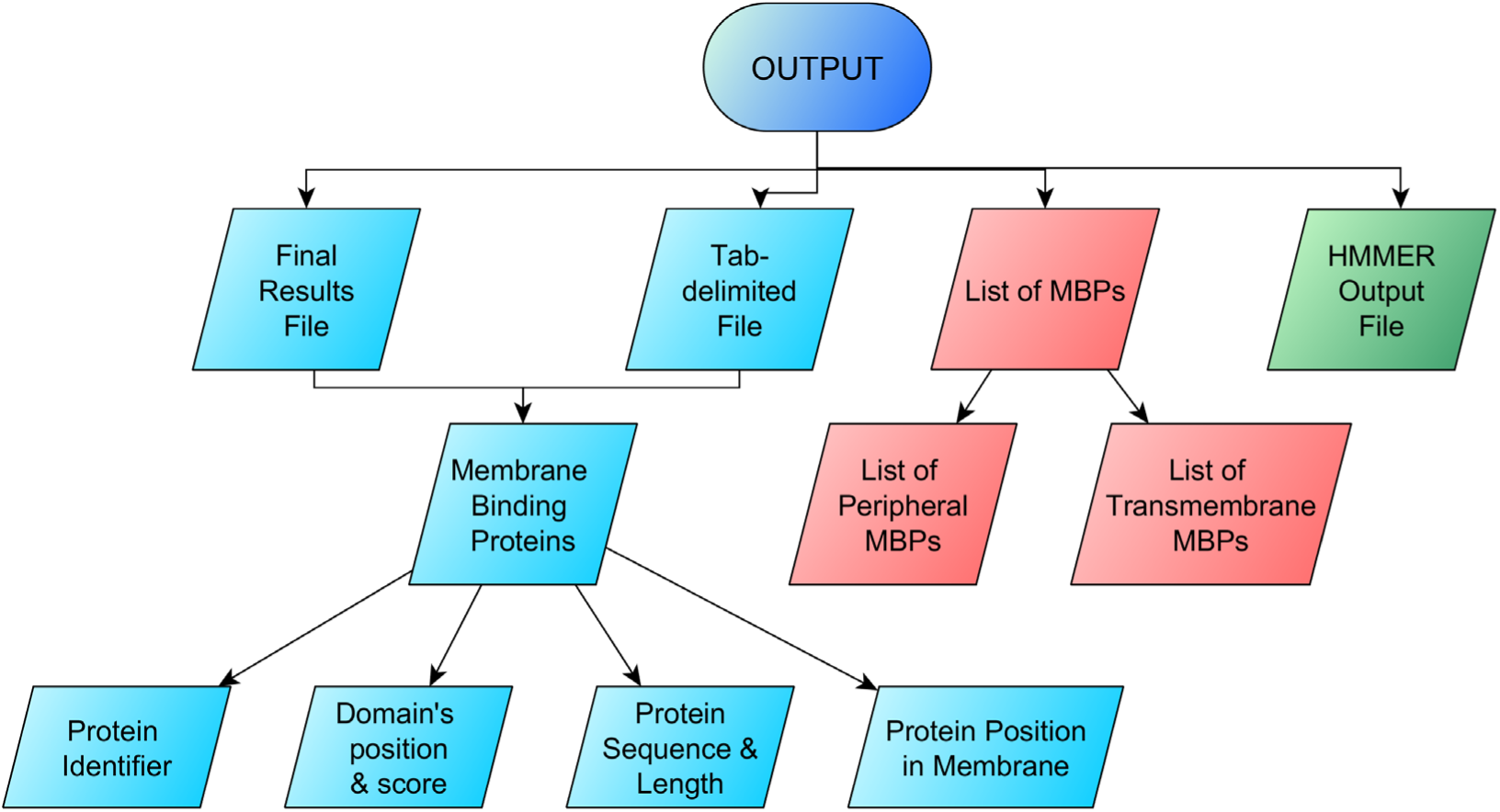
Output files produced by MBPpred and their contents

### 3.3. Evaluation of MBPpred

MBPpred performs very well during cross-validation, with high overall performance metrics (Table 2). The results from the jackknife test showed the method’s ability to correctly identify pseudo-novel sequences as MBPs (97.8% Sensitivity), while not erroneously detecting non-MBPs (99.9% Specificity).

**Table 2.**
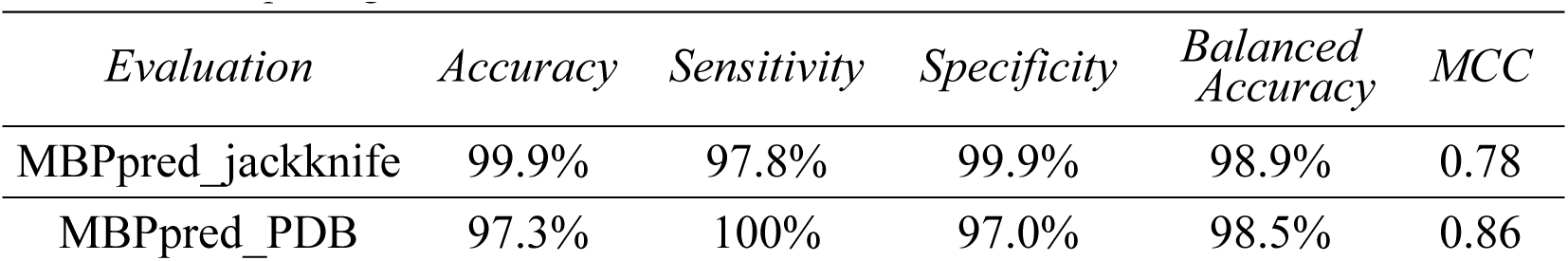
Results from the cross-validation of MBPpred using the jackknife technique and from the evaluation of MBPpred against the datasets assembled from PDB

| *Evaluation* | *Accuracy* | *Sensitivity* | *Specificity* | *Balanced Accuracy* | *MCC* |
| --- | --- | --- | --- | --- | --- |
| MBPpred_jackknife | 99.9% | 97.8% | 99.9% | 98.9% | 0.78 |
| MBPpred_PDB | 97.3% | 100% | 97.0% | 98.5% | 0.86 |

Our method was evaluated as a means to measure its performance against a non-redundant dataset of 49 membrane and 500 non-membrane binding proteins with known three-dimensional structure. Our method is accurate since it can detect all the proteins from the positive dataset as such (Sensitivity = 100%), while it falsely detects a very small percentage of non-MBPs as MBPs (Specificity = 97.0%) as shown in Table 2.

In addition, MBPpred was compared with the predictor developed by Bhardwaj et al. [17]. MBPpred outperforms this method, as shown in Table 3. We should note here that, our method could not be evaluated against the more recent method developed by Bhardwaj et al. [18], because the datasets used are not provided and none of the aforementioned methods are available online.

**Table 3.**
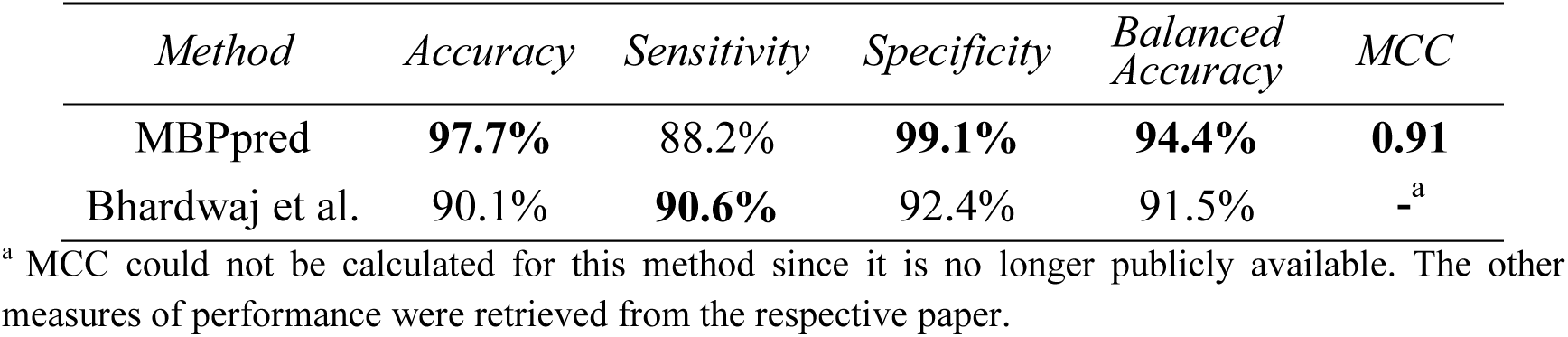
Comparison of MBPpred with the SVM method developed by Bhardwaj et al. (2006) ^a^ MCC could not be calculated for this method since it is no longer publicly available. The other measures of performance were retrieved from the respective paper.

### 3.4. Application of MBPpred in reference proteomes

The application of MBPpred in 30 eukaryotic reference proteomes showed that, ca. up to 6.0% of the proteins in these proteomes are possible MBPs (Fig. 3). The percentages vary based on the kingdom and phylum in which these organisms belong. In general, animals have more MBPs than fungi and plants, while other eukaryotes have a great divergence in the proportion of MBPs in their proteomes, whereas in general, organisms that are evolutionary closer to plants have less MBPs than organisms closer to animals and fungi.

**Figure 3.**
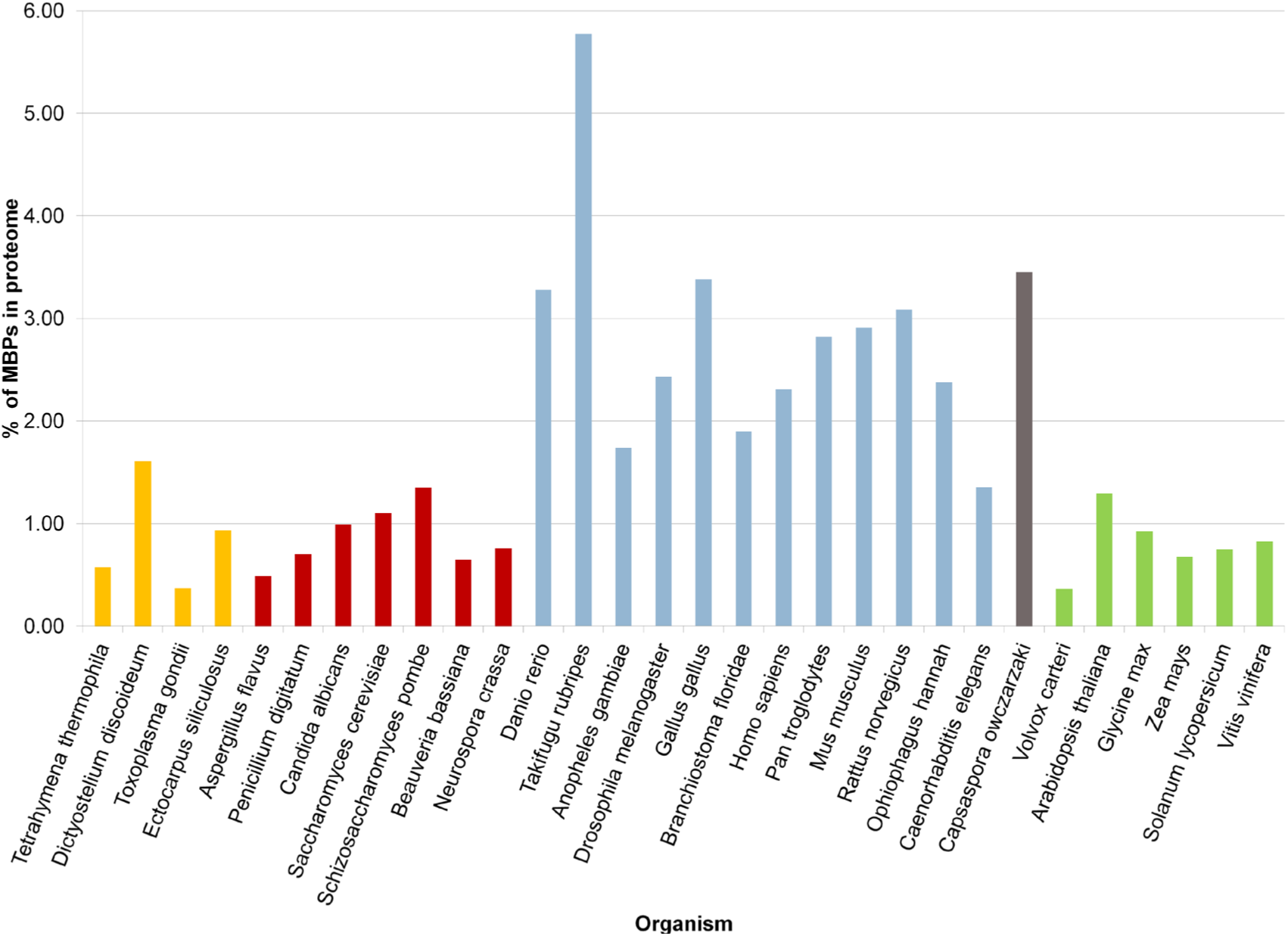
The percentage of MBPs in each of the 30 eukaryotic proteomes where MBPpred was applied. In general, Fungi (shown in red) and Plantae (shown in green) have less than 1.5% MBPs in their proteomes, while Animalia (light blue) have more than 1.5%. Organisms that belong to other eukaryotic kingdoms, like Amoebozoa, (orange and grey) have varying percentages of MBPs in their proteomes.

An enrichment analysis of the MBPs of 20 out of the 30 proteomes was performed (Table S5) using the DAVID functional annotation tool [26], in order to assess the functions of these proteins. Functional enrichment analysis could not be performed for MBPs from 10 proteomes (*Dictyostelium discoideum, Ectocarpus siliculosus, Candida albicans, Penicillium digitatum, Beauveria bassiana, Ophiophagus hannah, Capsaspora owczarzaki, Volvox carteri, Glycine max* and *Solanum lycopersicum*) because these proteomes have not been annotated with gene ontology terms. In all cases, terms related to lipid binding and certain membrane binding domains are overrepresented, as expected. Moreover, other terms associated with membrane trafficking and signal transduction are enriched, indicating the importance of MBPs in these cell processes. In particular, Gene Ontology (GO) [27] terms like regulation of Ras, Rho, small GTPase mediated and ARF protein signal transduction, protein kinase activity, endocytosis, cell junction and cytoskeleton organization are overrepresented in MBPs. There is no particular pattern in enriched terms – in any of the 3 major eukaryotic kingdoms – that would help us explain the differences in the percentages of MBPs between the studied organisms.

The classification of MBPs in peripheral membrane and transmembrane proteins, showed that in all cases peripheral MBPs are more than transmembrane. The deviation of the percentages in various kingdoms regarding MBPs can be attributed to the evolutionary diversity of these organisms (Fig. 4). Small differences in the membrane lipid and protein composition between these eukaryotes can be the cause of variability in the number of MBPs. Moreover, the big difference between animals and other eukaryotes can be attributed to the cell membrane differences of plants, fungi and animals, which consequently leads to differences in membrane protein composition [28]. Plants and fungi use different mechanisms to perform similar functions, like some of the functions in which MBPs take part in, e.g. endocytosis [29] and signal transduction [30–32] MBDs have been mainly studied in animals, and in particular mammals, and so, it is expected that evolutionary distant organisms will (or at list seem to) have less MBPs than those mainly studied with experimental and computational methods.

**Figure 4.**
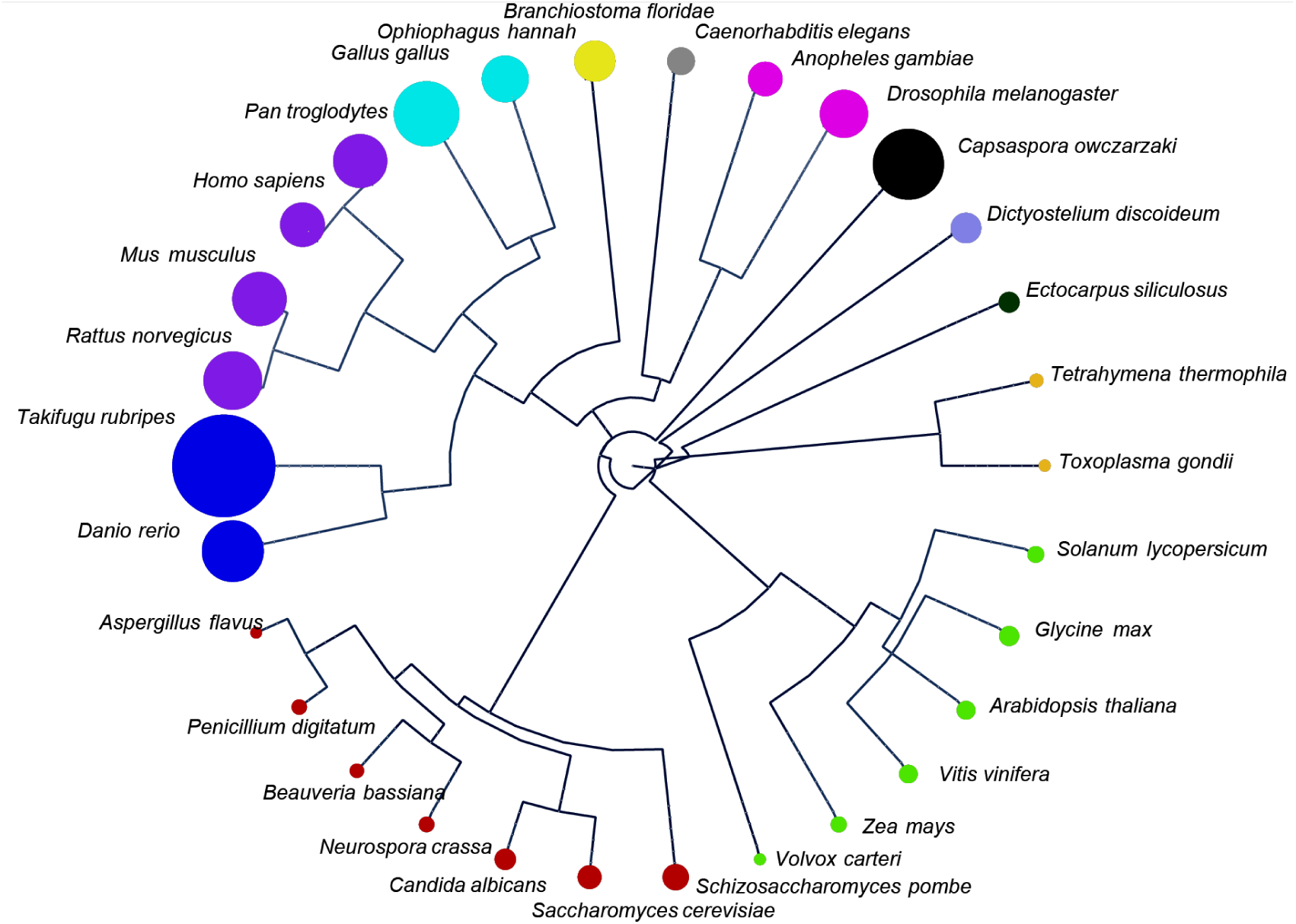
Cladogram for 30 selected proteomes. Each circle in the cladogram represents the percentage of MBPs in a proteome (from 0.3% in *Toxoplasma gondii* up to 5.7% in *Takifugu rubripes*). Each clade in the cladogram is color coded, where green represents plants, dark red represents fungi, dark blue is for fish, purple for mammals, light blue represents reptiles (including birds), yellow, grey and magenta other animals, orange amoebozoa and black, light purple and dark green represents organisms in other eukaryotic kingdoms. The cladogram was visualized with Cytoscape [33].

The classification of MBPs from the 30 eukaryotic proteomes in which MBPpred was applied showed that in peripheral membrane and transmembrane ca. 80% of MBPs are peripheral, while almost 20% are transmembrane (Table S6). Furthermore, there are some distinct differences regarding the MBD families present in various organisms (TableS7). While some of the most extensively studied MBDs (e.g. C2 and PH) are present in MBPs from all eukaryotic kingdoms, the more recently identified MBDs (e.g. Gla, PTB and IMD) are prevalently present in animals. The percentage of MBPs though, is generally similar within the different groups of eukaryotic proteomes. Interestingly, it was observed that almost 90% of all MBPs found in these 30 proteomes are proteins of unknown function or subcellular location. This was also observed during the functional annotation of the proteomes, where the majority of proteins could not be connected with any term. These results show that the application of MBPpred can greatly contribute to the understanding of proteins with unknown function, and, additionally help in the annotation of current and newly sequenced proteomes.

In addition to the large scale study that was performed, we used MBPpred to examine two well known human peripheral membrane binding proteins AKT1 and PTEN with great medical and clinical significance [34–38].

AKT1 is a RAC-α serine/threonine-protein kinase, which is involved in the regulation of many processes including metabolism, proliferation, cell survival, growth and angiogenesis [39]. AKT1 contains one pleckstrin homology (PH) domain, associated with the binding of membrane phosphoinositides [40]. Recent studies have shown that mutations of that domain in AKT1 are associated with various forms of cancer [34, 41, 42], and can enhance or impair the lipid binding properties of this protein [16]. After a literature search we were able to identify seven mutations (K8R, K14A, K14Q, K14R, K20Q, R25A, R86A) that are associated with reduced binding of various phosphoinositides, and one (E17K) which leads to enhanced binding [43–45]. We examined if MBPpred could detect that these proteins retain the ability to interact with the membrane, by identifying them as such, by applying MBPpred to all mutated AKT1 sequences. MBPpred successfully identified AKT1 as an MBP after substituting the amino acids connected with the mutations and re-submitting the protein sequence to the program. Further examination showed that the change of the domain’s score after each run of the method were consistent with the differences observed experimentally in binding affinity. Interestingly, in all 7 cases where the binding was impaired, the score was lower after the mutation, and in the case of the enhancing transforming mutation the score was higher.

PTEN is a phosphatidylinositol triphosphate phosphatase which acts as a tumor suppressor by negatively regulating Akt/PKB signaling pathway and is mutated in a large number of cancers [46, 47]. PTEN contains a C2 domain which plays a central role in membrane binding. Mutations of this domain have been associated with reduction in growth suppression activity and binding to phospholipid membranes [48]. In the case of PTEN we examined five mutations associated with the protein’s ability to retain its membrane binding (Y68H, R130L, R130Q, K289E, D331G) and two mutations associated with reduced membrane binding affinity (263-269 from KMLKKDK to AAGAADA and 327-335 from KANKDANR to AAGADAANA). In all cases MBPpred correctly identified the mutated protein sequences as membrane binding. The scores in the cases where the membrane binding affinity was retained were very close or exactly the same with those of the wild type protein after the application of MBPpred, while a reduction in the protein’s score was observed in the two mutations with experimentally identified impairment of membrane binding.

Consequently, the use of MBPpred in single protein sequences can also help in the better design of experiments regarding their lipid binding properties and aid towards the study of the mechanisms these proteins use to bind membranes.

MBPpred was also applied in all eukaryotic reference proteomes from UniProtKB (release: 2015_12) with similar results (Table S3). Additionally, in a search for homologous prokaryotic sequences MBPpred was applied in all 2629 Bacterial and 131 Achaean reference proteomes from UniProtKB (release: 2015_12) (TableS8). The description and study of MBDs in prokaryotes could help reveal the evolutionary origin for these protein domain families. Recent studies regarding the PH superfamilies have been conducted [49, 50] and MBPpred could aid immensely in such studies.

From the application of MBPpred in prokaryotic proteomes we observed that only 12 archaean and 3027 bacterial proteins are homologous to eukaryotic MBPs. From all 131 archaean reference proteomes only 4 contained at least one MBP-homolog. All 4 organisms (TableS8) belong to the Thaumarchaeota and Euryarchaeota phyla, which based on the eocyte theory are considered to be direct ancestors of eukaryotes [51, 52]. The analysis of the bacterial proteomes showed that 784 out of 2629 Bacterial proteomes contained at least one MBP-homolog, and there was no significant difference regarding the phyla that contained MBPs and those that did not. Moreover, there was no bacterial proteome that contained more than 1% of MBP homologs. A very interesting finding from the application of MBPpred in prokaryotic proteomes was that ~90% of the proteins identified by MBPpred contain the Discoidin domain, the only domain incorporated in MBPpred which has been previously identified in many bacterial proteins [53]. From the rest of the bacterial homologs ~8% contain the GOLPH3 domain, the most recently identified MBD in eukaryotes, whose function is currently being thoroughly studied and remains mainly unknown, and with less representatives there are proteins that contain Annexin, BAR, C1, C2, FYVE, GRAM and IMD domains.

The inability to identify proteins that interact with the membrane via MBDs in prokaryotes could be attributed to different mechanisms that these organisms use to mediate membrane binding and in the existence of analogous structural and functional characteristics they could possess that aid them towards performing these biological roles. Homologous domains to MBDs have been identified in prokaryotes – like bacterial domains that belong in the PH superfamily [10] – but the function of these domains still remains obscure.

In an effort to detect distant relationships between the 40 pHMMs incorporated in MBPpred and other domain families with representatives in Pfam we investigated the results that profile-profile alignments with the use of HHsearch [54] produced, in addition to the results of profile-profile comparisons using SCOOP [55]. From a total of 58 pHMMs identified in Pfam that are distantly related, 23 are already included in the pHMM library of MBPpred, showing that sequence similarities between MBDs exist, and that related domains perform indeed similar functions. The majority of the 35 domains that were found to be related to MBDs have very different well documented functions and five of them are domains of unknown function (DUFs) found in eukaryotes as well as prokaryotes (Table S9). Despite the fact that most of the domains with well characterized function do not perform lipid or membrane binding, they are in some way related to proteins or domains that perform these functions, either directly or indirectly. Additionally, many of these domains are zinc fingers mainly associated with C1, a zinc finger membrane binding domain that interacts with lipid substrates. The other zinc finger domains are also associated with binding, but of other substrates like DNA and RNA. Two very interesting cases were those of PhnJ and ACBP, both domains mainly found in prokaryotes [56, 57]. PhnJ is a phosphonate utilization domain [58], and even though this process is not relevant to membrane binding directly, it has been shown that phosphonate can act as an inhibitor to phosphate binding [59], showing that this distant relationship found in this case could help explain the divergence of function in related domains between prokaryotes and eukaryotes. Proteins that contain the ACBP domain have been found to interact with other large membrane associated proteins [57]. This indicates that although distant related protein families cannot be characterized as membrane binding, similar functions may be attributed to these protein families that would help in their experimental and computational study, in addition in the elucidation of their function.

## 4. CONCLUSIONS

MBPpred is a relatively fast and accurate method, which can detect Membrane Binding Proteins from their sequence alone and is therefore applicable to entire proteomes. Our method is the first to include an extended list of MBDs, compiled after an extensive literature search, for the detection of MBPs. Moreover, MBPpred can distinguish between peripheral and transmembrane MBPs and thus can identify peripheral membrane proteins, a group of proteins extremely challenging to predict and study from sequence alone [18]. In addition, MBPpred is currently the only publicly available method which can detect MBPs.

Even though experimental studies have shown that the overwhelming majority of proteins with Membrane Binding Domains have the ability to bind to phosphoinositides or other membrane lipids [60], there have been reports of a small number of proteins with MBDs, that have lost their ability to bind to membranes during the course of evolution [61]. However, the lack of experimental information regarding these proteins does not allow their discrimination from MBPs with the same domains. Nevertheless, their identification is crucial for their further functional annotation.

Computational studies for membrane binding proteins in organisms other than mammals have not been performed to date, and information gathered from the application of MBPpred on novel proteomes, can be of great assistance towards their functional annotation. The use of MBPpred for the annotation of newly sequenced proteomes is very important, since it can provide novel candidates for biochemical and structural analysis. Lipidomic studies have shown that cell membranes contain over 1000 different lipids [62]. Several of these lipids act as targets for Membrane Binding Proteins, which are recruited during cell signaling and membrane trafficking to form various protein-protein and lipid-protein interactions [2, 25]. These interactions are vital for the conduction of other membrane protein functions, since other membrane proteins with which MBPs interact can act as receptors, transporters, enzymes, structural proteins and so on [63]. In addition, a previous study of the peripheral membrane protein interactome (peripherome) of the human plasma membrane [64] indicated the importance of MBPs, since these proteins can act as hubs and bottlenecks in the network, while they maintain connections with other membrane proteins in microdomains that are enriched in certain membrane lipids, called lipid rafts [65]. The application employed here in the 30 and 407 proteome datasets is the first large scale effort for the identification of MBPs and provides important information regarding the presence and types of MBPs in various eukaryotic proteomes. Important insights are also gathered from the application of MBPpred in two selected proteins implicated in disease, AKT1 and PTEN, and also from the search of homologous sequences in prokaryotes. Utilizing profile-profile alignments [54] and profile-profile comparisons [55] for all 40 pHMMs against Pfam we were able to study remote homologies among eukaryotes and prokaryotes for MBDs and a list of putative domains involved directly or indirectly in membrane lipid binding was assembled. However, further studies are needed to safely associate these domains with membrane binding.

As more experimental information about MBPs becomes available, more proteins with the ability to bind to membrane lipids non-covalently will be revealed in all eukaryotic kingdoms. Consequently, more detailed information about the mechanism these proteins use to bind lipids will be uncovered and thus we will be able to better comprehend the interactions of proteins in the membrane plane.

## ACKNOWLEDGEMENTS

The authors would like to thank the scientific and administrative staff of the “Bioinformatics” Master’s Program at the Faculty of Biology of the University of Athens for generous support. The authors would also like to thank the anonymous reviewers and the handling editor for their valuable comments and constructive criticism.

## ABBREVIATIONS

*MBD(s):*: Membrane Binding Domain(s)
*MBP(s):*: Membrane Binding Protein(s)
*pHMM(s):*: profile Hidden Markov Model(s)
*MCC:*: Matthew’s Correlation Coefficient
*MSA:*: Multiple Sequence Alignment

